# Temperature influences immune cell development and body length in purple sea urchin larvae

**DOI:** 10.1101/2024.04.30.591967

**Authors:** Emily M. Wilkins, Audrey M. Anderson, Katherine M. Buckley, Marie E. Strader

## Abstract

Anthropogenic climate change has increased the frequency and intensity of marine heatwaves that may broadly impact the health of marine invertebrates. Rising ocean temperatures lead to increases in disease prevalence in marine organisms; it is therefore critical to understand how marine heatwaves impact immune system development. The purple sea urchin (*Strongylocentrotus purpuratus)* is an ecologically important, broadcast-spawning, omnivore that primarily inhabits kelp forests in the northeastern Pacific Ocean. The *S. purputatus* lifecycle includes a relatively long-lived (∼2 months) planktotrophic larval stage. Larvae have a well-characterized cellular immune system that is mediated, in part, by a subset of mesenchymal cells known as pigment cells. To assess the role of environmental temperature on the development of larval immune cells, embryos were generated from adult sea urchins conditioned at 14 °C. Embryos were then cultured in either ambient (14 °C) or elevated (18 °C) seawater. Results indicate find that larvae raised in an elevated temperature were slightly larger and had more pigment cells than those raised at ambient temperature. Further, the larval phenotypes varied significantly among genetic crosses, which highlights the importance of genotype in structuring how the immune system develops in the context of the environment. Overall, these results suggest that developmental temperature shapes the larval immune system and may adversely affect survival long-term.

## Introduction

Marine heatwaves (MHWs) are increasing in frequency and intensity due to global change and are characterized by periods of elevated sea surface temperatures that last for weeks to months and can span thousands of kilometers (Frölicher et al., 2018; Hobday et al., 2016). MHW events can have catastrophic impacts on marine habitats (*e.g.,* coral reef ecosystems (Fordyce et al., 2019) and kelp forests (Smale et al., 2019) and can have drastic impacts on marine communities through species range shifts (Sanford et al., 2019) decreased productivity (Whitney, 2015), altered food webs (Smith et al., 2021), and mass mortality events (Hoegh-Guldberg & Bruno, 2010; Laufkötter et al., 2020). MHW events are particularly stressful to benthic marine organisms, which are often unable to relocate to more favorable environments. As a result, during MHW events, benthic marine organisms must either have mechanisms to acclimate to their new environment via phenotypic plasticity or face potential mortality (Snell-Rood et al., 2018; West-Eberhard, 2003).

The increased prevalence of MHWs has magnified disease prevalence in the ocean (Burge et al., 2014; Rubio-Portillo et al., 2015). Elevated ocean temperatures can threaten biodiversity and survivability of marine organisms by affecting disease transmission, host susceptibility, pathogen survival and development (Harvell et al., 2002). There is also a positive correlation between growth rates of marine bacteria and fungi and increased temperature (Harvell et al., 2002). For example, increased temperature leads to a higher susceptibility of white band disease in reef-building corals (Bruno et al., 2007; Burge et al., 2014; Heron et al., 2010). The incidence of white band disease in *Acropora palmata* increases when median sea surface temperatures are ≥28.5 °C (Randall & Van Woesik, 2015). Although a limited number of mechanistic studies have been performed, two hypotheses for these observations have been proposed: 1) warmer waters relax the over-wintering dormancy of pathogenic microbes; and 2) persistent heat induces stress responses that suppress immune system function (Randall & Van Woesik, 2015). High temperatures have been implicated in the recent increased disease prevalence of sea star wasting disease (SSWD), which affects at least twenty species of sea stars off the west coast of North America (Bates et al., 2009; Eisenlord et al., 2016; Kohl et al., 2016; Miner et al., 2018). This disease has the potential to drastically impact community composition through local extinction events (Montecino-Latorre et al., 2016) and has led to trophic cascades resulting in kelp barrens and altered population structures (Schultz et al., 2016). In cooler water temperatures, SSWD progression slows but still results in mortality events, indicating that if elevated temperatures do subside, SSWD infections persist (Kohl et al., 2016). Since MHWs are projected to increase in intensity and severity over the coming years (Frölicher et al., 2018; Hobday et al., 2016), disease prevalence and host susceptibility will continue to plague marine organisms leading to potentially irreversible damage in marine ecosystems. Here, we examine the consequences of marine heatwaves on immune system development in another echinoderm species: the purple sea urchin (*Stronglyocentrotus purpuratus*).

*Stronglyocentrotus purpuratus* is an ecologically and economically important omnivore that inhabits the California Current System, which stretches from Baja California, Mexico to British Columbia, Canada (Checkley & Barth, 2009; Manier & Palumbi, 2008; Pearse, 2006). *S. purpuratus* are broadcast spawners with a biphasic life cycle that includes a long-lived planktotrophic larval stage that enables larvae to travel hundreds of kilometers on ocean currents (Okamoto et al., 2020; Pearse, 2006). Because pelagic *S. purpuratus* larvae may experience drastically different temperatures and environmental conditions to those of their parents, larvae exhibit the capacity to acclimate to variable environments (Gray, 2013; Leach & Hofmann, 2023; Puisay et al., 2018; Strader et al., 2022). It has been shown that elevated temperatures increase both growth and development rate in sea urchin larvae (Fujisawa & Shigei, 1990; O’Connor et al., 2007; Wong & Hofmann, 2020). Furthermore, temperatures experienced during early development has been shown to greatly influence survival in various tropical and temperate sea urchin species (Byrne et al., 2009; O’Connor et al., 2007; Sewell & Young, 1999). Therefore, it is important to understand if and how temperatures experienced during early embryogenesis influence the development of the immune system, which may subsequently affect survival later in ontogeny.

The purple sea urchin has a sophisticated and complex innate immune system (Smith, 2012) composed of several specialized cell types that mediate pathogen responses in both the adult and larval life stages (Rast et al., 2006; Smith et al., 2006). In *S. purpuratus*, larval immune cells are derived from mesenchymal cells that are specified in early embryogenesis during mid-blastula stage (Solek et al., 2013). These include pigment cells, which, in uninfected animals, exhibit a stellate morphology and primarily localize to the larval ectoderm, with concentrations in the tips of the arms and the apical end of the larvae (Ho et al., 2016; Smith et al., 2008). However, when exposed to certain strains of bacteria, pigment cells become active, change shape, and migrate to the site of infection (Ho et al., 2016).

Here, we identified how variation in developmental temperature impacts the morphology and immune cell development of *S. purpuratus* larvae. Specifically, we examined how developmental temperature and genotype impacted larval body size and immune system development by quantifying pigment cells. We find that larvae reared in elevated temperature were larger and had more pigment cells than those reared in ambient temperature. Furthermore, analysis of the individual genetic crosses revealed that genotype influences pigment cell count, but not overall body length. These results suggest that *S. purpuratus* larvae exhibit phenotypic plasticity in response to developmental temperature that shapes not only overall morphology, but also immune system development. Since marine heatwaves are not projected to cease in duration or intensity in the near future (Frölicher et al., 2018; Hobday et al., 2016), and coincide with increases in marine diseases, (Burge et al., 2014; Rubio-Portillo et al., 2015) these results highlight that phenotypic plasticity in immune cell development may enable *S. purpuratus* larvae to persist during periods of prolonged heat stress.

## Methods

### Conditioning of adult urchins

Adult *S. purpuratus* were collected off the coast of Santa Barbara, California by SCUBA in October 2021 (SBC LTER permit = California Department of Fish and Wildlife Scientific Collecting permit #SC-9228) and were transported to a saltwater tank facility at Auburn University, where animals were housed in an 85-gallon aquarium. Animals were maintained in artificial seawater (Instant Ocean; 14 °C; salinity = 30 ppt). Temperature and salinity were monitored daily using an Apollo IV DT304 Digital Temperature Logger (UEI) and an ATC refractometer; water chemistry was tested weekly using respective API test kits (API Saltwater Aquarium Master Test Kit). Adults were fed excess frozen kelp (*Macrocystis pyrifera*) once a week; a 20% water change occurred two days after feeding to stabilize water chemistry. Adults were acclimated to these conditions for three months prior to spawning.

### Spawning of adults and culturing of larvae

Adults were selected at random for spawning, which occurred in two rounds. Spawning was induced by an injection of 0.53 M KCl into the coelomic cavity. Sperm was collected dry and remained on ice until activation. Eggs were collected in 0.22 µm-filtered artificial sea water (FASW) at 30 ppt salinity and 14 °C temperature. Gamete compatibility was assessed by ensuring that fertilization success was >90%. For the first round of spawning, individual crosses were generated using one dam (Dam 1) and two sires (Sire 1 and 2). For the second round of spawning (which occurred two weeks later), individual crosses were established using a different set of adults, with two dams (Dam 2 and Dam 3) and one sire (Sire 3). Each spawning resulted in two unique genetic crosses, (4 total) from three males and three females. Fertilization occurred in ambient (14°C) FASW. Fertilized embryos from each cross were divided and cultured at one of temperatures: ambient (14 °C) and elevated (18 °C), each with two replicate cultures (Figure 1). Embryos were cultured in 4-liter vessels of FASW with stirring rotors at 20 rpm and a density of 10 embryos/µL (approximately 30,000 embryos per culture vessel).

**Figure 1:**
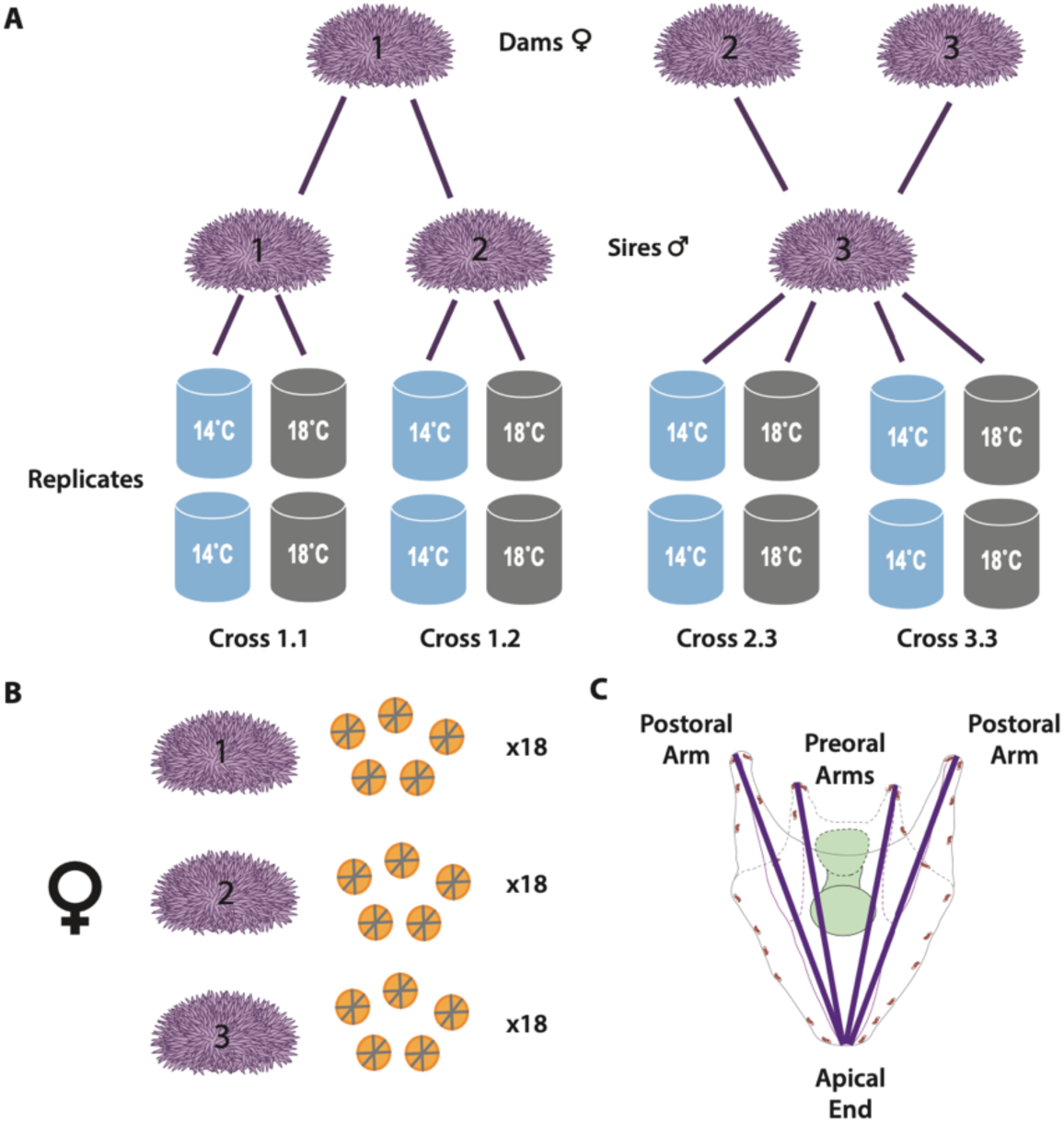
Experimental design used to investigate the role of developmental temperature on larval immune phenotypes. **(A)** Gametes were collected from three adult sires and three adult dams and crossed as shown. Cultures of fertilized embryos were divided and grown in either ambient (14 °C, shown in blue) or elevated (18 °C, shown in grey) temperatures. Each culture condition was performed as two replicates. (**B)** Morphological analysis was performed on unfertilized eggs from each dam (Figure 2). **(C)** Pluteus samples (6 dpf) were measured from each genetic cross and developmental condition. Preoral and postoral body length was measured (denoted with purple lines) and pigment cells (red) were counted.

### Early life-history sampling

Larval cultures were maintained for six days. Partial (1/3 volume) water changes were performed at 3 days post fertilization (dpf) to maintain water quality. Offspring were collected at pluteus stage (6 dpf) to quantify variation in body length and pigment cell count. Approximately 600 larvae from each culture vessel were preserved in 10% aqueous buffered zinc formalin (Z-Fix; Anatech, Ltd.), and stored at 4 °C prior to morphological analysis.

### Egg and early embryo morphology imaging and analysis

Unfertilized eggs from each female were also preserved in Z-Fix (approximately 600 eggs per female, collected in triplicate). Individual eggs (n=30 per female per triplicate) were selected at random and photographed using a Canon Rebel X digital camera and calibrated with a scale micrometer. Images were processed in FIJI (Schindelin et al., 2012). Average egg diameter was determined by taking the average of three independent diameter measurements at angles 0°, 45°, and 90° to account for potential irregularities in egg shape. To avoid bias in slide preps containing multiple eggs, each egg was randomly assigned a number and a random number generator was used to choose the egg to be measured.

### Pluteus morphology imaging and analysis

To measure pigment cell number and overall body length, larvae (N=10 per cross replicate) were imaged on a Zeiss Axio Observer 7 microscope with Zen Imaging Software (Zen 3.0 blue edition). Each image consisted of 50 slices of an interval of 0.57 µm. Images were processed using FIJI (Schindelin et al., 2012) using the “Cell Counter” plugin (Supplemental Figure 1). Pigment cells were counted manually. To accurately measure larval morphology, the X, Y, and Z coordinates were collected from each pre- and post-oral arm, as well as the apical end. The formula 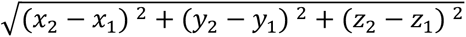 was used to calculate larval body length. Pre- and post-oral arm length was averaged for each individual larva.

### Statistical Analysis

All statistical analysis was conducted in R (version 4.1.2). Tests were run with a linear mixed effect model using the R data packages lme4 (Linear Mixed-Effects Models using ‘Eigen’ and S4, version 1.1-34; (Bates et al., 2015) and afex (Analysis of Factorial Experiments, version 1.3; (Singmann et al., 2023)). Models used to test if dam identity played a role in egg size, included fixed effects of dam and random effect of egg ID. To identify variation in pigment cell count, a linear mixed effect regression model (lmer) was used with developmental treatment and genetic combination used as fixed effects, while individual culture vessel used as a random effect. The interaction between genotype and treatment was a fixed effect in the model. Additional, separate lmer models were run to identify variation in pigment cell count due to maternal and paternal effects. The function emmeans (Estimated Marginal Means, aka Least-Squares Means, version 1.8.8, (Lenth, 2023)) was used to extrapolate individual comparisons. The same model structure was used to identify variation in preoral and postoral body length. Finally, a linear model using the lm package in R (lm: Fitting Linear Models, R stats package, version 4.2.3) was used to identify correlations between larval body length and pigment cell counts with developmental temperature used as the fixed effect. Separate correlations were performed to compare preoral/postoral body length and pigment cell count. Significance was defined as p<0.05.

## Results

### Egg diameter varies among dams

Unfertilized eggs were collected from the three dams to quantify variation in egg size. Analysis of egg diameters reveals that the three dams produced eggs of significantly different sizes (Figure 2). Dam 1 had the largest egg diameter (average = 94.17 µm; SE = 0.869 µm), while eggs from dams 2 and 3 were slightly smaller (average egg diameters of 88.93 ± 0.876 µm 88.13 ± 0.869 µm respectively). There was a significant difference in the diameter of eggs from dams 1 and 2 (*p*_lmer_ = 0.0119, Figure 2, Supplemental Table 1, 3) and dams 1 and 3 (*p*_lmer_ = 0.0066, Figure 2, Supplemental Table 1, 3). However, there was no significant difference in egg diameter between dam 2 and dam 3 (*p*_lmer_ = 0.8317, Figure 2, Supplemental Table 1, 3). Notably, a difference of 6 µm in egg diameter corresponds to up to a 14% difference in egg volume.

**Figure 2:**
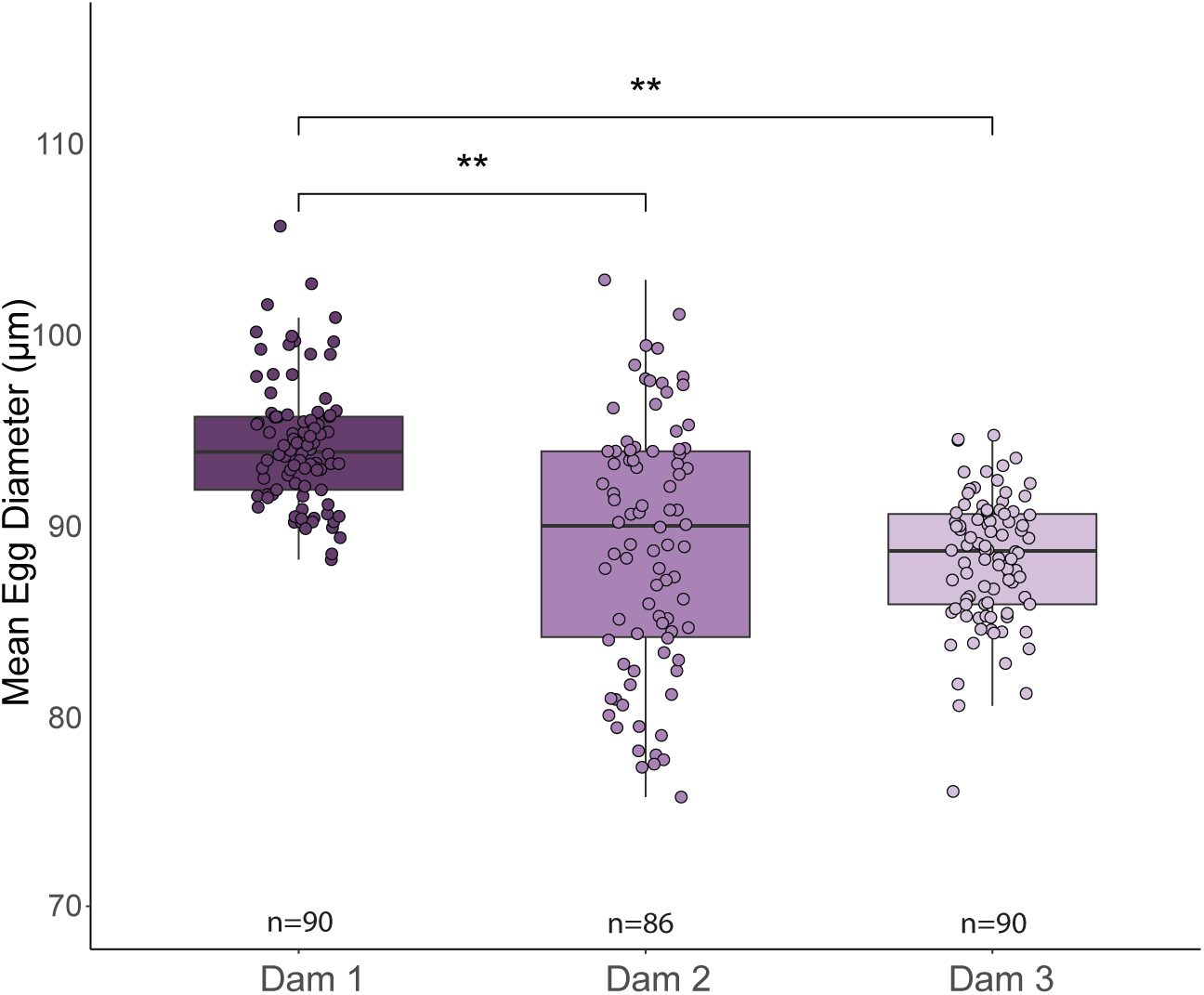
Egg diameter varies among dams. Unfertilized eggs were collected from each dam used in the experiment (n=90 per dam; indicated by shades of purple). Eggs were measured three times using orthogonal planes (0°, 45°, and 90°). The average of the three measurements from each egg is shown. Significant differences in egg size between individual dams are denoted with asterisks (*p*_lmer_ < 0.01).

### Larvae grow larger when cultured in higher temperatures

To determine the effects of genotype and temperature on preoral and postoral larval body length, we generated four genetic crosses from six adult sea urchins. Fertilized embryos were grown at either ambient (14 °C) or elevated (18 °C) temperatures (Figure 1). Larval morphology was measured at the 4-armed pluteus stage (6 dpf). At this stage, free-swimming larvae have fully developed gut structures and are able to feed. Additionally, pigment cells have terminally differentiated, ingressed through the blastocoel and are typically present near the ectoderm (Ho et al., 2016).

Postoral body length was significantly affected by temperature during embryogenesis. Pluteus larvae that developed in elevated temperatures were significantly larger than those grown in ambient conditions (*p*_lmer_ = 0.0324, Figure 3A). The average postoral body length of larvae cultured in elevated temperatures was an average of 276 µm (±14 µm, 95% CI); growing in ambient temperatures resulted in an average postoral body length of 253 µm (± 13.0 µm, 95% CI). We investigated potential influences of maternal or paternal effects on postoral body length and found that no significant differences independent of temperature conditions (Figure 3C, Supplemental Tables 14, 15; Figure 3D, Supplemental Tables 13, 15). Similar results were obtained from measurements of pre-oral body length (Supplementary Figure 2, Supplemental Tables 8-11).

**Figure 3:**
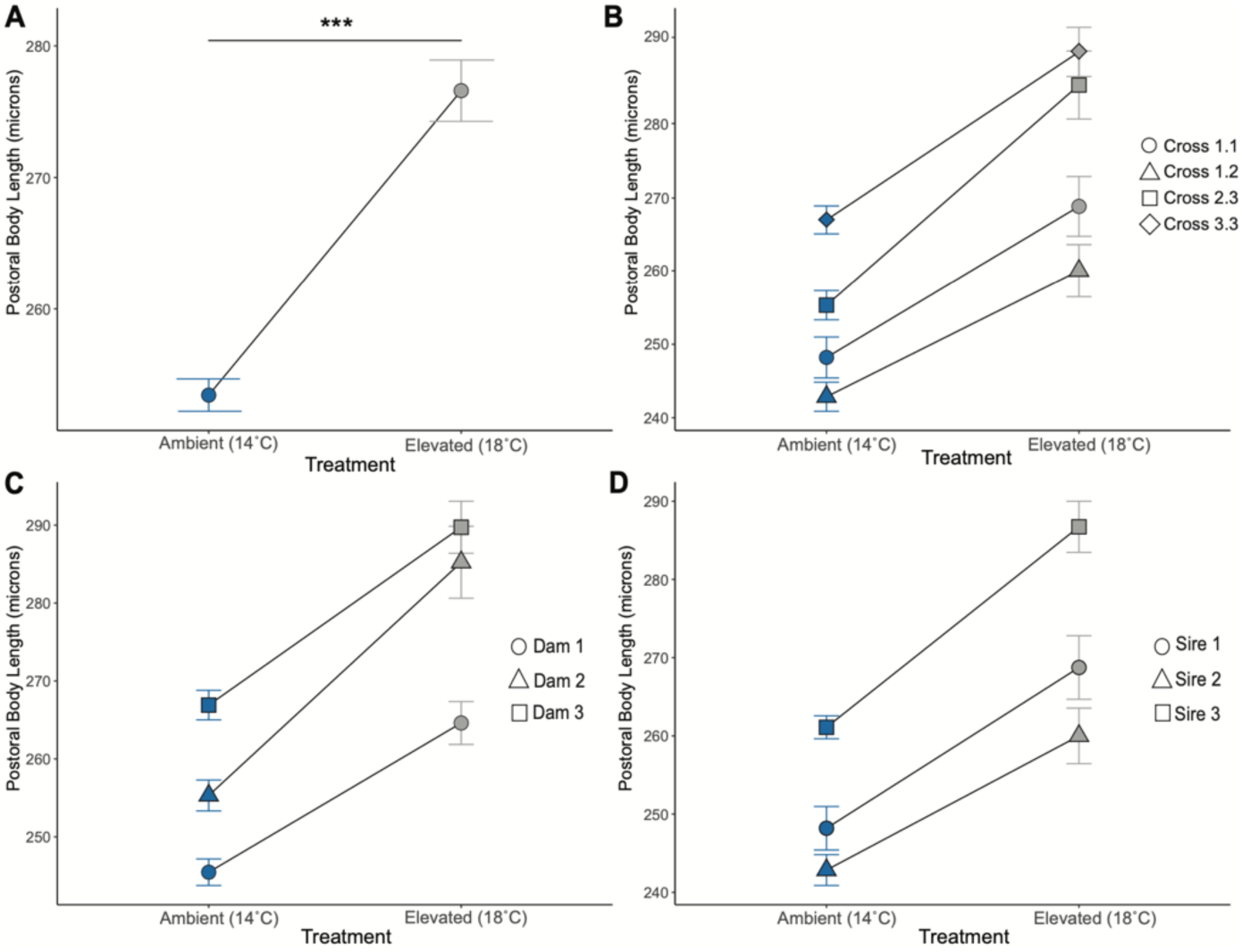
Embryos developed at elevated temperatures exhibit longer postoral body lengths. Individual adults used to generate the cultures are indicated by different shapes. Blue shapes indicate ambient conditions (14 °C); gray shapes indicate elevated conditions (18 °C). **(A)** Differences in postoral body length between larvae developed in ambient temperatures compared to larvae which developed in elevated temperatures (*p*_lmer_=0.0324). Asterisks denote significance differences (*p*_lmer_<0.001 = ***). No significant variation was observed in postoral body length differences based on genotype **(B)**, dams **(C)** or sires **(D)**.

### Higher temperatures during embryogenesis impact pigment cell development

In addition to identifying the effect of temperature and genotype on larval body length, we characterized how developmental temperature influences immune cell development. Given their importance in responding to immune challenge and distinctive morphology (Buckley & Rast, 2019; Ho et al., 2016), we enumerated pigment cells in each larva. On average, larvae reared in at 18 °C had more pigment cells than larvae grown in ambient conditions (*p*_lmer_ = 0.000431, Figure 4A).

**Figure 4:**
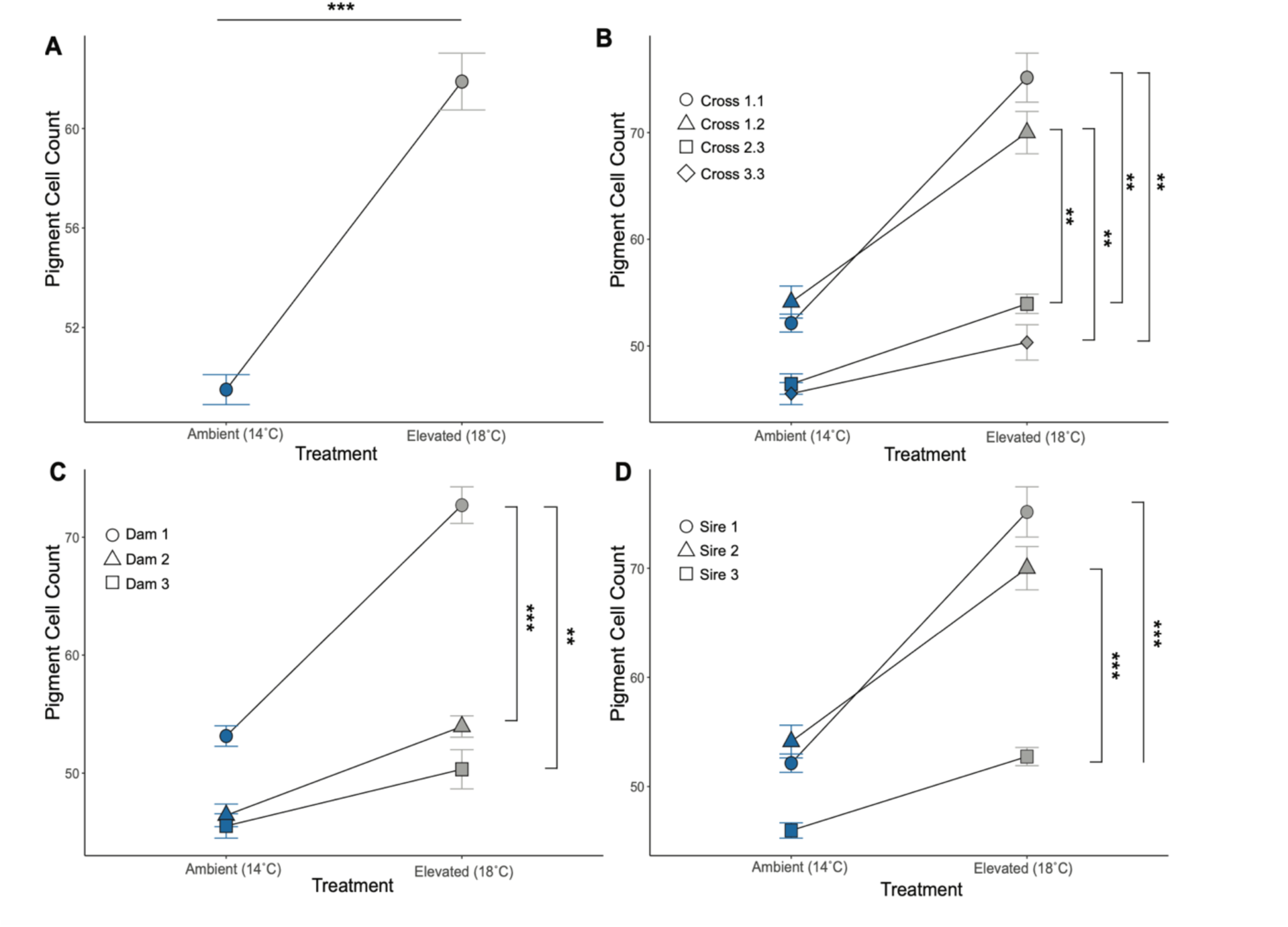
Pigment cell number is influenced by genotype and developmental temperature. **(A)** Differences in pigment cell count between larvae developed in ambient and elevated temperatures (*p_lm_* = 0.000431). **(B)** Differences in pigment cell count between unique genetic crosses. **(C)** Differences in pigment cell count between dams. **(D)** Differences in pigment cell count between sires. Asterisks denote significant differences (*p_lm_* <0.001 = ***; *p_lm_* <0.01 = **; *p_lm_* <0.05 =*).

We investigated the role of genotype, maternal and paternal effects on pigment cell development. Analyses of individual crosses revealed that genotype did not significantly influence pigment cell numbers for larvae reared in ambient temperature conditions but was an important factor in larvae reared in elevated temperatures (Figure 4B). Specifically, in elevated temperatures, larvae from cross 1.1 in had ∼25 more pigment cells than larvae from cross 3.3 (*p*_lmer_ = 0.0046), or cross 2.3 (*p*_lmer_ = 0.0034, Figure 4B, Supplemental Tables 4-5). However, the same crosses did not have significantly different pigment cell numbers under ambient conditions. Similarly, larvae from cross 1.2 had on average 16 more pigment cells when grown at 18 °C compared to the cross 2.3 (*p*_lmer_ = 0.0169) and ∼20 more pigment cells than cross 3.3 (*p_lmer_* = 0.0169, Figure 4B,Supplemental Tables 4-5).

To determine if maternal effects drive pigment cell variation among individuals, we compared larvae produced from the three dams. At ambient temperature, no significant variation was observed (Figure 4C, Supplemental Table 5). However, we found that dam played a significant role in pigment cell count in elevated temperatures. Under these conditions, larvae from dam 1 had on average 19 more pigment cells than larvae produced from dam 2 (*p*_lmer_ = 0.0009, Figure 4C, Supplementary Tables 5-6) or dam 3 (*p*_lmer_ = 0.0015, Figure 4C, Supplementary Tables 5-6). Additionally, there was no significant difference in pigment cell count between dam 2 and dam 3 under elevated temperature (*p_lm_* = 0.72, Figure 4C, Supplementary Table 5 & 6). Similarly to the maternal effects, we found that under ambient conditions, there were no significant differences between pigment cell count based on sire (Figure 4D, Supplementary Table 5 & 7), while effects were evident under elevated temperature. Larvae produced from sire 1 had on average 23 more pigment cells compared to larvae produced from sire 3 in under elevated temperature (*p*_lmer_ = 0.0001, Figure 4D, Supplementary Table 5 & 7). Additionally in the elevated temperature, larvae produced from sire 2 had significantly more pigment cells compared to larvae produced from sire 3 (*p*_lmer_ = 0.001, Figure 4D, Supplementary Table 5 & 7). Although there no significant different in pigment cell count between larvae produced from sire 1 and sire 2 in the elevated temperature, (*p*_lmer_ = 0.3454, Figure 4D, Supplementary Table 5 & 7).

Notably, there was no correlation between larval length and pigment cell number. Correlations between postoral body length and pigment cell count in ambient temperature, showed a negative, statistically significant, small correlation between the two variables (r^2^ = 0.041, 95% CI [-0.30, −0.05], *p* = 0.007). Additionally, under elevated temperature, we identified a negative, statistically not significant correlation between pigment cell count and postoral body length (r^2^ = 0.023, 95% CI [-0.30, 1.93e-03], *p* = 0.053).

Lastly, we quantified if egg size influenced larval postoral body length and pigment cell count for larvae in both developmental treatments. We found no significant correlation between egg diameter and postoral body length for larvae reared in ambient conditions for dam 1 (r^2^ = 0.0041, *p* = 0.629, Figure 6A), for dam 2 (r^2^ = 0.0029, *p* = 0.684, Figure 6A) or for dam 3 (r^2^ = 0.002, *p* = 0.734, Figure 6A), or for larvae reared in elevated conditions for dam 1 (r^2^ = 0.18, *p* = 0.333, Figure 6C, or for dam 2 (r^2^ =6.4 x 10^-6^, *p* = 0.989, Figure 6C). However, we did identify a positive, significant, correlation between egg diameter and postoral body length for dam 3 in elevated conditions (r^2^ = 0.21, *p* = 0.012, Figure 6C). Similarly, there was no significant correlation for samples reared in ambient conditions originating from dam 1 (r^2^ = 0.00099, *p* = 0.811) or from dam 2 (r^2^ = 0.0018, *p* = 0.746). However, there is a positive, statistically significant correlation between pigment cell count and egg diameter between samples reared in ambient conditions originating from dam 3 (r^2^ = 0.083, *p* = 0.025, Figure 6B). For samples reared in the elevated conditions, we found no significant correlation between egg diameter and pigment cell count between samples reared in elevated conditions originating from dam 1 (r^2^ = 0.0029, *p* = 0.779, Figure 6D), dam 2 (r^2^ = 0.0045, *p* = 0.725, Figure 6D), or dam 3 (r^2^ =3.9e-06, *p* = 0.992, Figure 6D).

**Figure 5:**
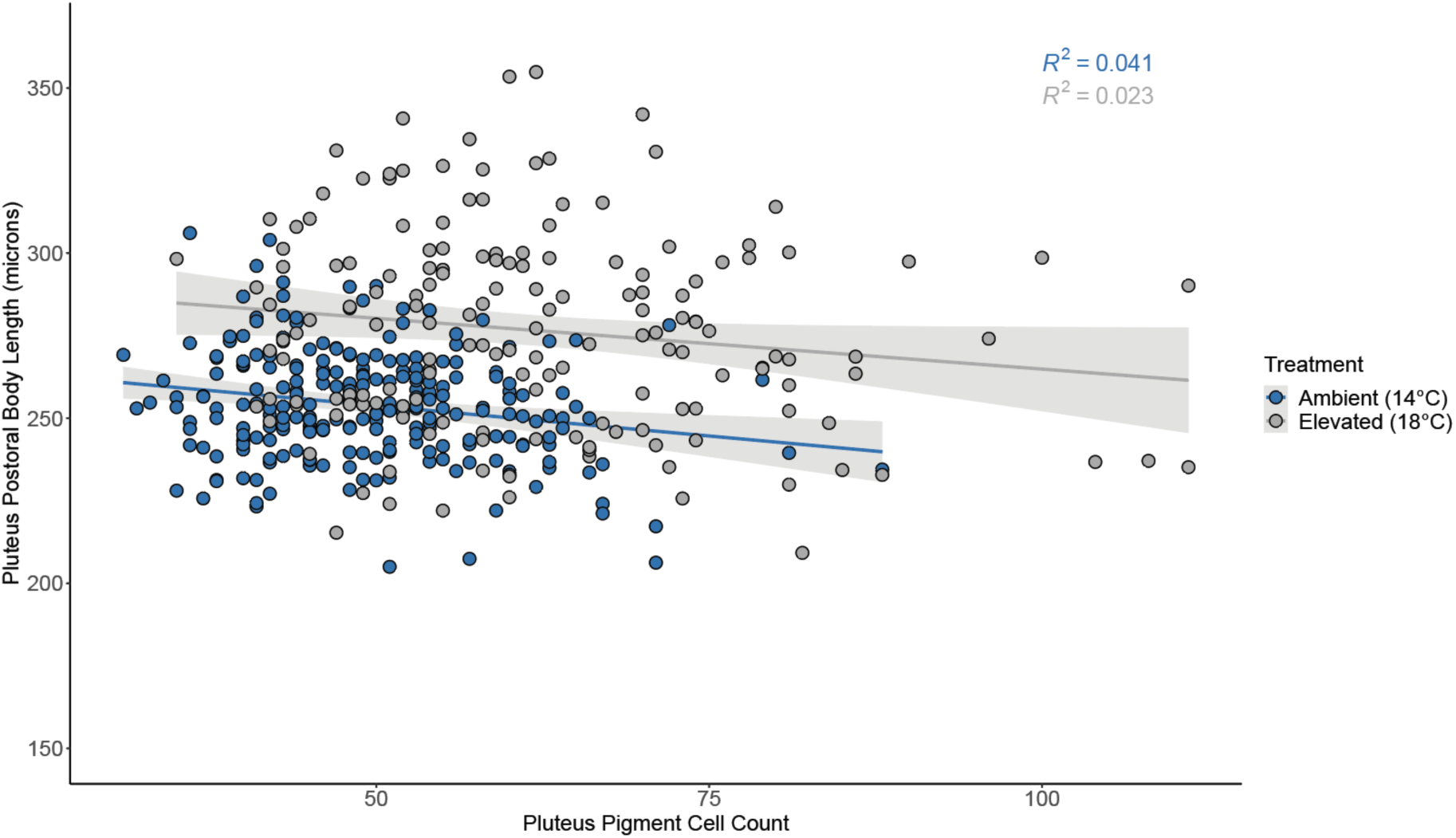
Relationship between pigment cell count and postoral body length. Blue dots represent larvae reared in ambient temperatures (14 °C) while gray dots represent larvae reared in elevated temperatures (18 °C). Grey shading indicates standard error.

**Figure 6:**
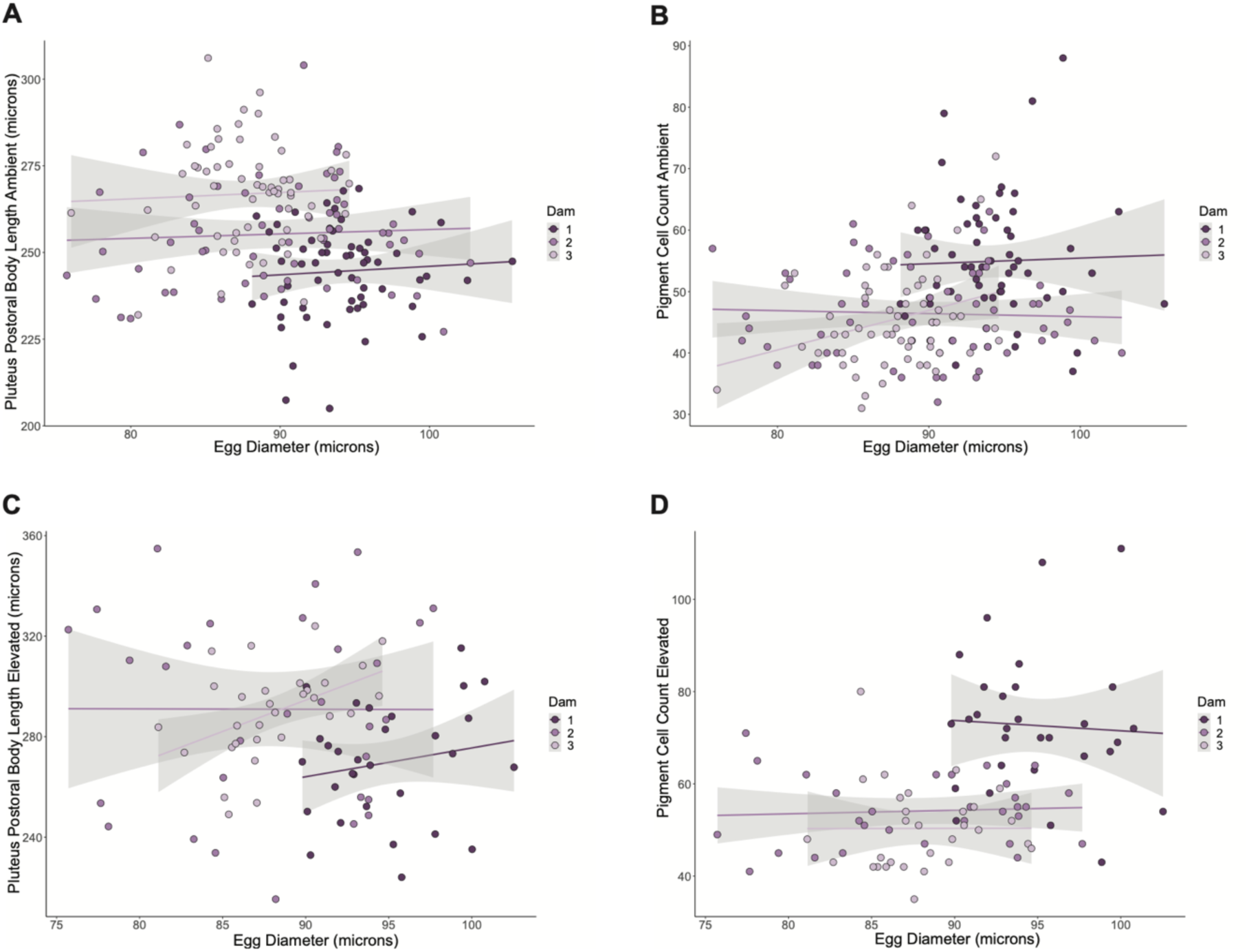
Relationship between egg diameter and postoral body length. **(A)** or pigment cell count **(B)** of larvae reared in ambient (14 °C) temperatures. Relationship between egg diameter and larval postoral body length **(C)** and pigment cell count **(D)** for larvae developed in elevated (18 °C) temperature. The three shades of purple represent the dams used in this experiment. Points represent individual larvae sampled, while gray shading represent standard error.

## Discussion

We investigated the role of temperature on body length and immune cell development in *S*. *purpuratus* larvae and found that elevated temperature during embryogenesis resulted in larvae that were larger and had more pigment cells. In addition to the environmental influence, this variation is also driven by genotype but not by maternal provisioning, as inferred through differences in egg size. Overall, these results suggest that environmental temperature is a primary driver of plasticity during development.

### Egg diameter varies among dams

In broadcast spawning marine organisms, egg size is one of the most important factors contributing not only to fertilization success, but to larval growth, development, and ultimately survival (Moran & McAlister, 2009). Although egg size is critical for species success, this phenotype is extremely variable not only among species, but also among individuals. Therefore, in the context of how early developmental environments shape phenotypes, it is important to have a comprehensive understanding of when variation occurs, and what might drive these differences. Here, we found significant variation in egg diameter between dams. However, when identifying if variation in egg size influences larval postoral body length or pigment cell count, we found that there was no correlation between egg size on those two factors, regardless of developmental condition. This suggests that maternal provisioning, which we infer from unfertilized egg size, was not a contributing factor shaping later larval size and immune capabilities.

Intraspecific variation in egg size is not uncommon in broadcasting spawning marine organisms (Emlet & Hoegh-Guldberg, 1997; Levitan, 2000, 2006; Marshall et al., 2000; McEdward & Carson, 1987) and can be driven by environmental conditions, genetic variation, or stochastic developmental processes (Moran & McAlister, 2009; Vogt et al., 2008). The temperature experienced by dams during oogenesis can influence the size of the egg produced (Moran & McAlister, 2009), but likely did not cause the sizes differences in the eggs observed here, as the urchins used for this experiment were collected during the same dive event at the same site and were housed in the same common garden aquarium for three-months prior to spawning. Additionally, adult urchins did not spawn during this three-month acclimation period. However, we are unaware of the conditions they experienced in the wild, and their previous spawning history in the wild, which would most likely have a stronger influence on their egg size and the intraspecific variation we observed in our study.

Maternal age can also influence the size of the egg produced, with egg size decreasing with advanced maternal age in marine invertebrates (Ito, 1997; Moran & McAlister, 2009; Qian & Chia, 1992). Since our adult urchins were collected from the wild, we are unable to know for certain their age and if this is contributing to the variation we observed in egg diameter. Maternal size can also contribute to overall egg size such that larger mothers produce larger eggs (George, 1994; Marshall et al., 2000; Moran & McAlister, 2009) and may reflect the overall amount of resources that can be directed to reproduction (George, 1994). However, our data display the opposite trend. Although sea urchin adults were randomly selected at the time of spawning, the test size of dam 1 was the smallest (diameter = 56 cm; height = 35 cm) and produced the largest eggs, while dam 3 was the largest (diameter = 73 cm; height 38 cm) and produced the smallest eggs (Supplemental Table 16). Thus, it is likely that additional factors other than only maternal size may contribute to the observation variation in egg diameters.

For broadcast-spawning marine invertebrates, the entire nutritive contribution from the mother to offspring is provided in the egg, which may result in larger eggs that are better equipped to handle developmental stressors (McEdward & Carson, 1987). Additionally, larger eggs have a great surface area to enable sperm interactions, which is extremely beneficial for broadcast-spawners, especially in sperm-limited conditions (Levitan, 1996; Marshall & Keough, 2007). However, in areas with high densities of males, the influence of egg size on survival may be reverse, as polyspermy occurs more frequently in large eggs (Marshall & Keough, 2007). Thus, egg size is likely subject to a variety of evolutionary pressures. After fertilization, egg size can greatly influence larval growth, development, and survival (Emlet, 1995; Levitan, 2000; Marshall & Keough, 2007).

### Developmental temperature influences body length in larval sea urchins

The environment in which an individual develops has monumental influence over their success and survival as an adult (Gray, 2013; Marshall et al., 2003; Shima & Findlay, 2002). When faced with adverse environmental conditions, organisms can either relocate to a more favorable site, adjust to their new environment, or perish. However, larval marine organisms are often subjected to the will of the currents, and therefore unable to relocate. Temperature strongly influences larval development; understanding how developmental temperature shapes larval phenotypes is extremely important in understanding how organisms will respond to changing environments (Byrne, 2011; Byrne & Przeslawski, 2013; O’Connor et al., 2007; Pechenik, 1984). Here, we show that pluteus cultured in elevated temperatures were significantly larger than those grown in ambient temperatures. These results indicate that larval sea urchins are phenotypically plastic to variation in temperature. The ability to produce a plastic response when exposed to changes in environmental condition is critical for organismal survival, particularly during sensitive developmental stages. However, this allocation of resources may come at a cost to developing larvae and have downstream consequences after metamorphosis to adulthood. Plasticity resulting from variation in developmental temperatures has been well-documented in marine invertebrate larvae (Byrne, 2011; Byrne & Przeslawski, 2013; González-Ortegón & Giménez, 2014; O’Connor et al., 2007; Reitzel et al., 2004), and, more specifically, in many echinoderms species (Byrne & Przeslawski, 2013; Dellatorre & Manahan, 2023; Hoegh-Guldberg & Pearse, 1995; Karelitz et al., 2020; Leach & Hofmann, 2023; Wangensteen et al., 2013; Wong & Hofmann, 2020). Our results add to this rich literature base.

The mechanistic driver that enables increased larval body length is an increase in metabolic activity (Byrne, 2011; Sardi et al., 2023; Somero, 2002). By increasing the rate of biochemical reactions, elevated temperatures can affect metabolic rates (Dellatorre & Manahan, 2023; Sin et al., 2019). In *S. purpuratus*, higher temperatures have been shown to increase not only metabolic rates, but also overall body length and arm length (Dellatorre & Manahan, 2023; Leach & Hofmann, 2023; Strader et al., 2022). Longer arms increase food acquisition rates, which may compensate for increased energy needs (Dellatorre & Manahan, 2023). Additionally, amino acid transporters in the larval arms of purple sea urchins are critical for metabolic processes such as osmoregulation, nutrition, and protein synthesis, and are in direct contact with seawater, to facilitate the transport of environmental nutrients (Christensen et al., 1965; Meyer & Manahan, 2009).

In addition to morphological and metabolic changes resulting from variation in temperature, there are ecological consequences of planktonic larvae developing in increased water temperatures. Increased developmental temperatures have been shown decrease the time to metamorphosis (Byrne, 2011; O’Connor et al., 2007). Shorter planktonic larval durations may benefit larvae by decreasing rates of predation and increasing settlement rates and overall survival (Allen, 2008; Byrne et al., 2011; Hare & Cowen, 1997). For instance, under elevated temperature conditions, planktonic larvae of *Rhopaloeides odorabile*, a common sponge found in the Great Barrier Reef, settled 36 hours faster than those at ambient temperature conditions (Whalan et al., 2008). Shortened planktonic larval duration can limit genetic connectivity between populations since larvae spend less time traversing the water column (O’Connor et al., 2007). Additionally, since predators prefer smaller larvae, increasing larval size may influence the predator-prey dynamics of marine food webs (Allen, 2008). Since marine heat waves (MHWs) are projected to increase in intensity and duration (Frölicher et al., 2018; Hobday et al., 2016), and SST are rising at alarming rates (Hoegh-Guldberg et al., 2014), our results reveal challenges that are likely to be faced by all larval marine organisms, including *S. purpuratus*.

Lastly, to identify whether genotype influences postoral body length during larval development, we generated four genetic crosses and exposed embryos from each cross to both experimental conditions until they reached 6 dpf (Figure 1). We found that genotype did not significantly impact postoral body length, regardless of developmental temperature. These results indicate developmental temperature is the primary driver of plasticity in larval length. Our results differ, however, from previous research investigating genotypic effects influencing larval morphometrics. In Leach & Hofmann, 2023, researchers found that both paternal identity, and the interaction between paternal identity and larval environment, significantly impacted larval arm length in *S. purpuratus* larvae. Although they investigated an earlier echinopluteus larval stage, these results indicate that there may be additional drivers influencing larval plasticity in stressful environments, or that genotypic effects on larval body morphometrics are prominent at earlier developmental stages and may cease in later larval life.

### Developmental temperature and genotype influence pigment cell count in larval sea urchins

Planktonic larvae are directly exposed to the open ocean and need to protect themselves against environmental pathogens to reach adulthood. However, changes in environmental conditions can lead to an increase in pathogens (Cohen et al., 2018; Vezzulli et al., 2013), thus necessitating a robust immune system. Here, we find that larvae exposed to increased temperatures have significantly more pigment cells than those grown in ambient temperatures. These results demonstrate that larval sea urchins can alter the development of their immune system after prolonged exposure to adverse environmental conditions, which may serve to better protect themselves from harmful pathogens present in their environment.

Pigment cells are unique to echinoid larvae, which makes this an interesting model to investigate environmental impacts on immune system development (Spurrell et al., 2023). Pigment cells are often used as an indicator for immune response in purple sea urchin larvae due to their red coloration, which makes cells easily quantifiable and traceable in transparent larvae. Further, there is a well characterized suite of genes expressed in pigment cells, including polyketide synthase 1 (*SpPks1*), flavin-dependent monooxygenase 3 (*SpFmo3*) and macrophage migration inhibitory factor 5 (*SpMif5*) (Spurrell et al., 2023). These pigment cell precursor genes are first expressed in the blastula stage and remain expressed throughout development (Spurrell et al., 2023). Although we know the genes responsible specifying pigment cells, and how these cells respond to pathogenic stressors, this is the first study to examine how these cells respond to altered environmental conditions. Future work should examine if the genes which regulate pigment cells production are modulated during embryogenesis in different environmental conditions to generate the increased pigment cell numbers observed here or if the pigment cells themselves divide in response to elevated temperatures.

While our study is the first to connect temperature with immune cell development in larval sea urchins, ocean temperatures have dramatic impacts on immune responses in other marine invertebrates. In prolonged exposure to elevated water temperatures, abalones (*Haliotis rubra)* exhibit compromised antibacterial responses and increases in antiviral activity, indicating a potential trade-off in immune response due to heat stress (Dang et al., 2012). Additionally, larval lobsters (*Homarus americanus*) grown in elevated temperature conditions have higher total hemocyte numbers which serves as an indicator of the innate immune response (Harrington et al., 2019). These results highlight the ability of marine organisms to adjust their immune system to better defend themselves against pathogens.

Similar to the *S. purpuratus* larval stage, the adult purple sea urchin has a complex immune system that responsive to environmental change and is mediated by a heterogeneous suite of coelomocytes (Barela Hudgell et al., 2022; Matranga et al., 2000, 2005; Rast et al., 2006; L. C. Smith et al., 2006). In response to temperature stress, coelomocytes increase in concentration (particularly red sphere cells and phagocytes) (Branco et al., 2012) and upregulate heat shock proteins (Matranga et al., 2000). In some cases, sea urchin coelomocytes are even used as bioindicators of environmental stress (Matranga et al., 2000). Thus, as in the larval sea urchins analyzed here, immune cell populations in adult sea urchins are capable of responding to environmental stressors. Larvae that have more immune cells may be able to more efficiently respond to pathogens. However, this may come at a cost: by allocating resources to increasing pigment cell numbers, other developmental processes may be compromised. Additionally, not much is known about how the quantity of pigment cells in larval stages influences coelomocyte concentration in adulthood, so this variation could have consequences in adult sea urchins as well.

Lastly, we found that genotype contributed significantly to variation in pigment cell count, but only for larvae developed at higher temperatures, suggesting genotype x environment (GxE) interactions. Similarly, in Pacific oyster (*Crassostrea gigas*) larvae, there is a significant interaction between environmental temperature, host genotype, and pathogen genotype (GxGxE), with disease resistance occurring in the warmer experimental conditions (Wendling et al., 2017). These results indicate that there is a strong genetic basis to the development of the immune system, and that larvae exposed to elevated temperature have the potential to be better-equipped to handle environmental pathogens. Larvae in our experiment were grown in sterile-filtered (0.2 micron) ASW and were not knowingly exposed to bacteria. Thus, the responses observed are likely based on factors controlled for in the experiment (i.e. temperature and genotype), and not as a response to additional pathogens.

## Conclusions

We find that *S. purpuratus* larvae are phenotypically plastic in response to developmental temperature. This plasticity enables larvae to modulate their total number of immune cells, which may affect their ability to defend against potential pathogens. Since marine heatwaves are projected to increase in duration and intensity in the near future (Frölicher et al., 2018; Hobday et al., 2016), coinciding with increases in marine diseases, (Burge et al., 2014; Rubio-Portillo et al., 2015) these results highlight that phenotypic plasticity in immune cell development may enable *S. purpuratus* larvae to persist during periods of prolonged heat stress.

## Data Availability

Raw data files and analysis scripts can be found at: https://github.com/emw0083/MorphologyData

## Acknowledgements

This work was supported by a National Science Foundation award to KMB and MES (2323328) as well as start-up funds provided by Auburn University. Additional support was provided by Auburn University’s CASE REU program for sponsoring AMA. The authors would like to thank Dr. Gretchen Hofmann for providing the animals used in this study, and Dr. Todd Steury and Dr. Dan Warner for statistical council. Additional thanks to Megan Maloney, Felicity Milam, and Kiley Ekert for assistance with egg imaging and measurements.

