## Supplemental Text for "Temperature influences immune cell development and body length in purple sea urchin larvae"

EMW: 0009-0006-8349-1109

KMB: 0000-0002-6585-8943

MES: 0000-0002-1886-4187

**Keywords:** *Strongylocentrotus purpuratus*; marine heatwaves; immune cell development; larval morphology

**Supplemental Table 1**: Spreadsheet containing egg diameter measurements for each of the three dams sampled. This spreadsheet is included as a separate excel file and can be found at: https://github.com/emw0083/MorphologyData.

**Supplemental Table 2:** Spreadsheet containing pigment cell counts and preoral and postoral body length measurements for all individuals sampled. This spreadsheet is included as a separate excel file and can be found at: https://github.com/emw0083/MorphologyData

| **Supplemental Table 3:** Egg size variability between the three dams sampled |  |  |  |
| --- | --- | --- | --- |
| **Test** | **df** | **t-value** | **p-value** |
| Dam 1 vs Dam 2 | 6.03 | 4.304 | **0.0119** |
| Dam 1 vs. Dam 3 | 5.94 | 4.909 | **0.0066** |
| Dam 2 vs. Dam 3 | 6.03 | 0.589 | 0.8317 |

**Supplemental Table 4:** Pigment cell mean values for genetic cross

| **Treatment** | **Cross** | **emmean** | **SE** | **df** | **lower CL** | **upper CL** |
| --- | --- | --- | --- | --- | --- | --- |
| Ambient (14˚C) | 1.1 | 52 | 2.66 | 6.71 | 45.7 | 58.4 |
| Ambient (14˚C) | 1.2 | 54.1 | 2.65 | 6.62 | 47.8 | 60.5 |
| Ambient (14˚C) | 2.3 | 46.6 | 2.65 | 6.57 | 40.1 | 52.8 |
| Ambient (14˚C) | 3.3 | 45.5 | 2.65 | 6.57 | 39.2 | 51.9 |
| Elevated (18˚C) | 1.1 | 75.2 | 2.78 | 7.95 | 68.7 | 81.6 |
| Elevated (18˚C) | 1.2 | 69.8 | 2.83 | 8.47 | 63.3 | 76.2 |
| Elevated (18˚C) | 2.3 | 54 | 2.65 | 6.57 | 47.6 | 60.3 |
| Elevated (18˚C) | 3.3 | 50.3 | 3.75 | 6.57 | 41.4 | 59.3 |

**Supplemental Table 5:** Significance values of pigment cell counts

| **Treatment** | **Test** | **SE** | **df** | **t-value** | **p-value** |
| --- | --- | --- | --- | --- | --- |
| Ambient (14˚C) | Cross 1.1 vs Cross 1.2 | 3.76 | 6.66 | -0.556 | 0.9417 |
| Ambient (14˚C) | Cross 1.1 vs Cross 2.3 | 3.76 | 6.64 | 1.491 | 0.4918 |
| Ambient (14˚C) | Cross 1.1 vs Cross 3.3 | 3.76 | 6.64 | 1.731 | 0.3798 |
| Ambient (14˚C) | Cross 1.2 vs Cross 2.3 | 3.75 | 6.59 | 2.052 | 0.2607 |
| Ambient (14˚C) | Cross 1.2 vs Cross 3.3 | 3.75 | 6.59 | 2.292 | 0.1938 |
| Ambient (14˚C) | Cross 2.3 vs Cross 3.3 | 3.75 | 6.57 | 0.24 | 0.9946 |
| Ambient (14˚C) | Dam 1 vs Dam 2 | 3.25 | 8.4 | 2.047 | 0.1602 |
| Ambient (14˚C) | Dam 1 vs Dam 3 | 3.25 | 8.4 | 2.324 | 0.1067 |
| Ambient (14˚C) | Dam 2 vs Dam 3 | 3.75 | 8.36 | 0.24 | 0.9688 |
| Ambient (14˚C) | Sire 1 vs Sire 2 | 3.46 | 8.65 | -0.602 | 0.8227 |
| Ambient (14˚C) | Sire 1 vs Sire 3 | 3 | 8.65 | 2.02 | 0.165 |
| Ambient (14˚C) | Sire 2 vs Sire 3 | 2.99 | 8.55 | 2.723 | 0.0578 |
| Elevated (18˚C) | Cross 1.1 vs Cross 1.2 | 3.96 | 8.21 | 1.362 | 0.5531 |
| Elevated (18˚C) | Cross 1.1 vs Cross 2.3 | 3.84 | 7.24 | 5.521 | **0.0034** |
| Elevated (18˚C) | Cross 1.1 vs Cross 3.3 | 4.67 | 7.02 | 5.319 | **0.0046** |
| Elevated (18˚C) | Cross 1.2 vs Cross 2.3 | 3.87 | 7.49 | 4.08 | **0.0169** |
| Elevated (18˚C) | Cross 1.2 vs Cross 3.3 | 4.69 | 7.18 | 4.138 | **0.0169** |
| Elevated (18˚C) | Cross 2.3 vs Cross 3.3 | 4.59 | 6.57 | 0.788 | 0.8576 |
| Elevated (18˚C) | Dam 1 vs Dam 2 | 3.31 | 9.02 | 5.604 | **0.0009** |
| Elevated (18˚C) | Dam 1 vs Dam 3 | 4.24 | 8.75 | 5.228 | **0.0015** |
| Elevated (18˚C) | Dam 2 vs Dam 3 | 4.59 | 8.36 | 0.788 | 0.72 |
| Elevated (18˚C) | Sire 1 vs Sire 2 | 3.68 | 11.03 | 1.461 | 0.3454 |
| Elevated (18˚C) | Sire 1 vs Sire 3 | 3.26 | 9.76 | 6.879 | **0.0001** |
| Elevated (18˚C) | Sire 2 vs Sire 3 | 3.3 | 10.21 | 5.165 | **0.001** |

**Supplemental Table 6:** Emmean for dam pigment cell count

| **Treatment** | **Dam** | **emmean** | **SE** | **df** | **lower CL** | **upper CL** |
| --- | --- | --- | --- | --- | --- | --- |
| Ambient (14˚C) | 1 | 53.1 | 1.88 | 8.48 | 48.8 | 57.4 |
| Ambient (14˚C) | 2 | 46.4 | 2.65 | 8.36 | 40.4 | 52.5 |
| Ambient (14˚C) | 3 | 45.5 | 2.65 | 8.36 | 39.5 | 51.6 |
| Elevated (18˚C) | 1 | 72.5 | 1.98 | 10.43 | 68.1 | 76.9 |
| Elevated (18˚C) | 2 | 54 | 2.65 | 8.36 | 47.9 | 60 |
| Elevated (18˚C) | 3 | 50.3 | 3.75 | 8.36 | 41.8 | 58.9 |

**Supplemental Table 7:** Emmean for sire pigment cell count

| **Treatment** | **Sire** | **emmean** | **SE** | **df** | **lower CL** | **upper CL** |
| --- | --- | --- | --- | --- | --- | --- |
| Ambient (14˚C) | 1 | 52 | 2.45 | 8.72 | 46.5 | 57.9 |
| Ambient (14˚C) | 2 | 54.1 | 2.44 | 8.57 | 48.6 | 59.7 |
| Ambient (14˚C) | 3 | 46 | 1.72 | 8.51 | 42 | 49.9 |
| Elevated (18˚C) | 1 | 75.2 | 2.58 | 10.64 | 69.5 | 80.8 |
| Elevated (18˚C) | 2 | 69.8 | 2.63 | 11.43 | 64 | 75.5 |
| Elevated (18˚C) | 3 | 52.7 | 1.99 | 8.51 | 48.2 | 57.3 |

**Supplemental Table 8:** Emmean for genetic cross preoral body length

| **Treatment** | **Genetic Cross** | **emmean** | **SE** | **df** | **lower CL** | **upper CL** |
| --- | --- | --- | --- | --- | --- | --- |
| Ambient (14˚C) | 1.1 | 246 | 11.6 | 6.96 | 219 | 274 |
| Ambient (14˚C) | 1.2 | 243 | 11.6 | 6.94 | 215 | 270 |
| Ambient (14˚C) | 2.3 | 262 | 11.6 | 6.93 | 235 | 290 |
| Ambient (14˚C) | 3.3 | 254 | 11.6 | 6.93 | 227 | 282 |
| Elevated (18˚C) | 1.1 | 272 | 11.7 | 7.13 | 245 | 300 |
| Elevated (18˚C) | 1.2 | 256 | 11.7 | 7.2 | 229 | 284 |
| Elevated (18˚C) | 2.3 | 287 | 11.6 | 6.93 | 259 | 314 |
| Elevated (18˚C) | 3.3 | 269 | 16.4 | 6.93 | 230 | 308 |

**Supplemental Table 9:** Emmean for dam preoral body length

| **Treatment** | **Dam** | **emmean** | **SE** | **df** | **lower CL** | **upper CL** |
| --- | --- | --- | --- | --- | --- | --- |
| Ambient (14˚C) | 1 | 244 | 7.73 | 8.91 | 227 | 262 |
| Ambient (14˚C) | 2 | 262 | 10.93 | 8.88 | 237 | 287 |
| Ambient (14˚C) | 3 | 254 | 10.93 | 8.88 | 229 | 279 |
| Elevated (18˚C) | 1 | 264 | 7.8 | 9.22 | 247 | 282 |
| Elevated (18˚C) | 2 | 287 | 1093 | 8.88 | 262 | 311 |
| Elevated (18˚C) | 3 | 269 | 15.45 | 8.88 | 234 | 304 |

**Supplemental Table 10:** Emmean for sire preoral body length

| **Treatment** | **Sire** | **emmean** | **SE** | **df** | **lower CL** | **upper CL** |
| --- | --- | --- | --- | --- | --- | --- |
| Ambient (14˚C) | 1 | 246 | 10.94 | 8.96 | 221 | 271 |
| Ambient (14˚C) | 2 | 243 | 10.93 | 8.93 | 218 | 268 |
| Ambient (14˚C) | 3 | 258 | 7.73 | 8.92 | 241 | 276 |
| Elevated (18˚C) | 1 | 272 | 11.01 | 9.21 | 248 | 297 |
| Elevated (18˚C) | 2 | 256 | 11.05 | 9.32 | 231 | 281 |
| Elevated (18˚C) | 3 | 281 | 8.92 | 8.92 | 261 | 301 |

**Supplemental Table 11:** Significance values for preoral body length values

| **Treatment** | **Test** | **SE** | **df** | **t-value** | **p-value** |
| --- | --- | --- | --- | --- | --- |
| Ambient (14˚C) | Cross 1.1 vs Cross 1.2 | 16.4 | 6.95 | 0.208 | 0.9965 |
| Ambient (14˚C) | Cross 1.1 vs Cross 2.3 | 16.4 | 6.94 | -0.98 | 0.7653 |
| Ambient (14˚C) | Cross 1.1 vs Cross 3.3 | 16.4 | 6.94 | -0.492 | 0.9584 |
| Ambient (14˚C) | Cross 1.2 vs Cross 2.3 | 16.4 | 6.93 | -1.188 | 0.6529 |
| Ambient (14˚C) | Cross 1.2 vs Cross 3.3 | 16.4 | 6.93 | -0.7 | 0.8939 |
| Ambient (14˚C) | Cross 2.3 vs Cross 3.3 | 16.4 | 6.93 | 0.488 | 0.9594 |
| Ambient (14˚C) | Dam 1 vs Dam 2 | 13.4 | 8.89 | -1.328 | 0.4163 |
| Ambient (14˚C) | Dam 1 vs Dam 3 | 13.4 | 8.89 | -0.731 | 0.7523 |
| Ambient (14˚C) | Dam 2 vs Dam 3 | 15.5 | 8.88 | 0.518 | 0.8649 |
| Ambient (14˚C) | Sire 1 vs Sire 2 | 15.5 | 8.95 | 0.22 | 0.9737 |
| Ambient (14˚C) | Sire 1 vs Sire 3 | 13.4 | 8.95 | -0.901 | 0.653 |
| Ambient (14˚C) | Sire 2 vs Sire 3 | 13.4 | 8.93 | -1.156 | 0.5064 |
| Elevated (18˚C) | Cross 1.1 vs Cross 1.2 | 16.5 | 7.17 | 0.985 | 0.7624 |
| Elevated (18˚C) | Cross 1.1 vs Cross 2.3 | 16.5 | 7.03 | -0.864 | 0.8231 |
| Elevated (18˚C) | Cross 1.1 vs Cross 3.3 | 20.1 | 6.99 | 0.166 | 0.9982 |
| Elevated (18˚C) | Cross 1.2 vs Cross 2.3 | 16.5 | 7.07 | -1.851 | 0.3268 |
| Elevated (18˚C) | Cross 1.2 vs Cross 3.3 | 20.1 | 7.02 | -0.642 | 0.915 |
| Elevated (18˚C) | Cross 2.3 vs Cross 3.3 | 20.1 | 6.93 | 0.874 | 0.8181 |
| Elevated (18˚C) | Dam 1 vs Dam 2 | 13.4 | 8.99 | -1.663 | 0.2705 |
| Elevated (18˚C) | Dam 1 vs Dam 3 | 17.3 | 8.95 | -0.276 | 0.9592 |
| Elevated (18˚C) | Dam 2 vs Dam 3 | 18.9 | 8.88 | 0.928 | 0.6376 |
| Elevated (18˚C) | Sire 1 vs Sire 2 | 15.6 | 9.26 | 1.043 | 0.5695 |
| Elevated (18˚C) | Sire 1 vs Sire 3 | 14.2 | 9.09 | -0.59 | 0.8288 |
| Elevated (18˚C) | Sire 2 vs Sire 3 | 14.2 | 9.16 | -1.735 | 0.2443 |

**Supplemental Table 12:** Emmeans for postoral body length for genetic cross

| **Treatment** | **Genetic Cross** | **emmean** | **SE** | **df** | **lower CL** | **upper CL** |
| --- | --- | --- | --- | --- | --- | --- |
| Ambient (14˚C) | 1.1 | 248 | 14.2 | 6.96 | 215 | 282 |
| Ambient (14˚C) | 1.2 | 243 | 14.2 | 6.94 | 209 | 76 |
| Ambient (14˚C) | 2.3 | 255 | 14.2 | 6.94 | 222 | 289 |
| Ambient (14˚C) | 3.3 | 267 | 14.2 | 6.94 | 233 | 301 |
| Elevated (18˚C) | 1.1 | 269 | 14.3 | 7.11 | 235 | 302 |
| Elevated (18˚C) | 1.2 | 259 | 14.3 | 7.17 | 225 | 292 |
| Elevated (18˚C) | 2.3 | 285 | 14.2 | 6.94 | 252 | 319 |
| Elevated (18˚C) | 3.3 | 290 | 20.1 | 6.94 | 242 | 337 |

**Supplemental Table 13:** Emmean for sire postoral body length

| **Treatment** | **Sire** | **emmean** | **SE** | **df** | **lower CL** | **upper CL** |
| --- | --- | --- | --- | --- | --- | --- |
| Ambient (14˚C) | 1 | 248 | 12.87 | 8.96 | 219 | 278 |
| Ambient (14˚C) | 2 | 243 | 12.86 | 8.93 | 214 | 272 |
| Ambient (14˚C) | 3 | 261 | 9.09 | 8.92 | 241 | 282 |
| Elevated (18˚C) | 1 | 269 | 12.95 | 9.2 | 240 | 298 |
| Elevated (18˚C) | 2 | 259 | 12.99 | 9.3 | 230 | 288 |
| Elevated (18˚C) | 3 | 287 | 10.5 | 8.92 | 263 | 311 |

**Supplemental Table 14:** Emmeans for dam for postoral body length

| **Treatment** | **Dam** | **emmean** | **SE** | **df** | **lower CL** | **upper CL** |
| --- | --- | --- | --- | --- | --- | --- |
| Ambient (14˚C) | 1 | 246 | 9.07 | 8.91 | 225 | 266 |
| Ambient (14˚C) | 2 | 255 | 12.82 | 8.89 | 226 | 284 |
| Ambient (14˚C) | 3 | 267 | 12.82 | 8.89 | 238 | 296 |
| Elevated (18˚C) | 1 | 264 | 9.15 | 9.21 | 243 | 284 |
| Elevated (18˚C) | 2 | 285 | 12.82 | 8.89 | 256 | 314 |
| Elevated (18˚C) | 3 | 290 | 18.13 | 8.89 | 249 | 331 |

**Supplemental Table 15:** Significance values for postoral body length

| **Treatment** | **Test** | **SE** | **df** | **t-value** | **p-value** |
| --- | --- | --- | --- | --- | --- |
| Ambient (14˚C) | Cross 1.1 vs Cross 1.2 | 20.1 | 6.95 | 0.279 | 0.9918 |
| Ambient (14˚C) | Cross 1.1 vs Cross 2.3 | 20.1 | 6.95 | -0.344 | 0.9848 |
| Ambient (14˚C) | Cross 1.1 vs Cross 3.3 | 20.1 | 6.95 | -0.921 | 0.7953 |
| Ambient (14˚C) | Cross 1.2 vs Cross 2.3 | 20.1 | 6.94 | -0.623 | 0.9215 |
| Ambient (14˚C) | Cross 1.2 vs Cross 3.3 | 20.1 | 6.94 | -1.2 | 0.6462 |
| Ambient (14˚C) | Cross 2.3 vs Cross 3.3 | 20.1 | 6.94 | -0.577 | 0.9359 |
| Ambient (14˚C) | Dam 1 vs Dam 2 | 15.7 | 8.89 | -0.62 | 0.8134 |
| Ambient (14˚C) | Dam 1 vs Dam 3 | 15.7 | 8.89 | -1.358 | 0.4014 |
| Ambient (14˚C) | Dam 2 vs Dam 3 | 18.1 | 8.89 | -0.64 | 0.8026 |
| Ambient (14˚C) | Sire 1 vs Sire 2 | 18.2 | 8.95 | 0.308 | 0.9493 |
| Ambient (14˚C) | Sire 1 vs Sire 3 | 15.8 | 8.95 | -0.808 | 0.7079 |
| Ambient (14˚C) | Sire 2 vs Sire 3 | 15.7 | 8.93 | -1.164 | 0.5021 |
| Elevated (18˚C) | Cross 1.1 vs Cross 1.2 | 20.3 | 7.14 | 0.494 | 0.9579 |
| Elevated (18˚C) | Cross 1.1 vs Cross 2.3 | 20.2 | 7.02 | -0.817 | 0.8448 |
| Elevated (18˚C) | Cross 1.1 vs Cross 3.3 | 24.7 | 7 | -0.85 | 0.8296 |
| Elevated (18˚C) | Cross 1.2 vs Cross 2.3 | 20.2 | 7.06 | -1.312 | 0.5842 |
| Elevated (18˚C) | Cross 1.2 vs Cross 3.3 | 24.7 | 7.02 | -1.255 | 0.6157 |
| Elevated (18˚C) | Cross 2.3 vs Cross 3.3 | 24.6 | 6.94 | -0.182 | 0.9976 |
| Elevated (18˚C) | Dam 1 vs Dam 2 | 15.8 | 8.99 | -1.363 | 0.3985 |
| Elevated (18˚C) | Dam 1 vs Dam 3 | 20.3 | 8.95 | -1.279 | 0.441 |
| Elevated (18˚C) | Dam 2 vs Dam 3 | 22.2 | 8.89 | -0.202 | 0.9777 |
| Elevated (18˚C) | Sire 1 vs Sire 2 | 18.3 | 9.25 | 0.545 | 0.8513 |
| Elevated (18˚C) | Sire 1 vs Sire 3 | 16.7 | 9.09 | -1.079 | 0.5494 |
| Elevated (18˚C) | Sire 2 vs Sire 3 | 16.7 | 9.15 | -1.676 | 0.2651 |

**Supplemental Table 16:** Morphological measurements of the six adults used in this experiment.

| **Individual** | **Diameter** | **Height** |
| --- | --- | --- |
| Sire 1 | 72 cm | 37 cm |
| Sire 2 | 56 cm | 33 cm |
| Sire 3 | 64 cm | 34 cm |
| Dam 1 | 56 cm | 35 cm |
| Dam 2 | 70 cm | 40 cm |
| Dam 3 | 73 cm | 38 cm |

### **Supplemental Results**

### *Larvae grow larger when cultured in higher temperatures*

Results indicate that preoral body length was significantly affected by temperature during embryogenesis. Pluteus larvae that developed in elevated temperatures were significantly larger (average length of 272 ± 12.0 µm, 95% CI; n = 166) than larvae grown in ambient temperatures (average length of 252 ± 11.0 µm, 95% CI; n = 235; *p*_lmer_=0.0211, Figure S2A). To investigate the possibility that maternal effects influence preoral body length, we identified variation in overall preoral body length in larvae produced by the three dams. Independent of temperature conditions, we found that there was no significant difference in preoral body length based on dam (Figure S2C, Supplemental Tables 9, 11). To determine if paternal effects drive preoral body length, we quantified preoral body length based on sire. We found that there were no significant paternal effects on preoral body length, independent of temperature treatment (Figure S2D, Supplemental Tables 10, 11). Finally, no significant genotypic effect was observed on preoral body length when comparing our four genetic crosses (Supplemental Tables 8, 11; Figure S2B).

**Supplemental Figures**

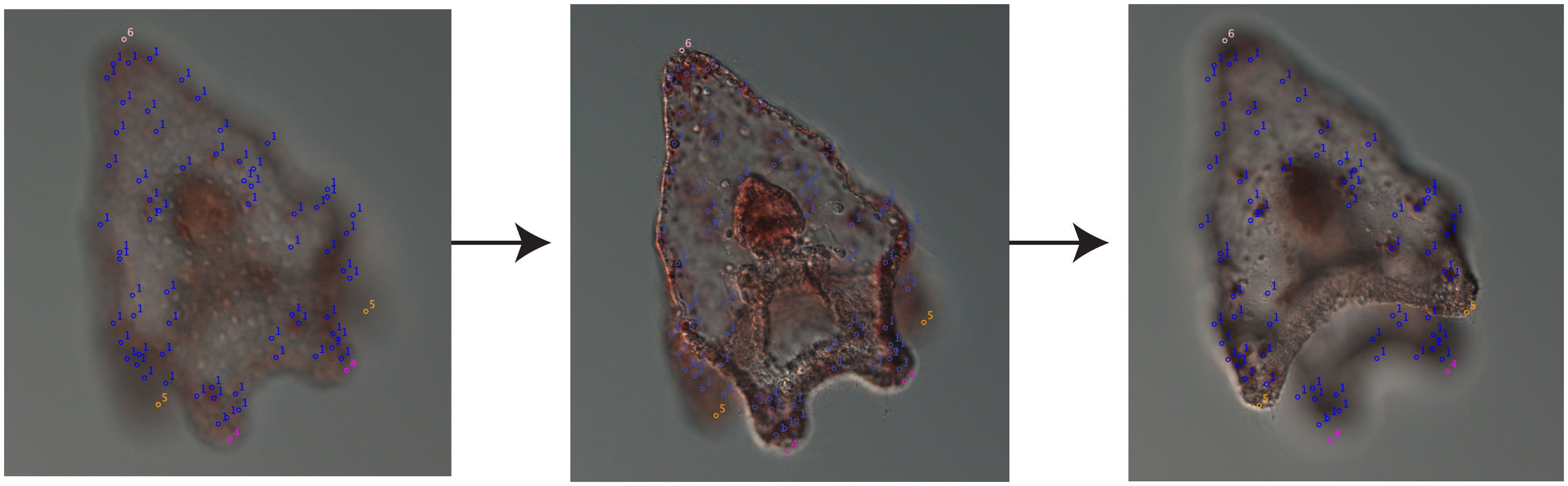

**Supplemental Figure 1:** Quantifying pigment cell count and preoral and postoral body length in pluteus larvae. A total of 50 slices per individual Z-stack were taken to quantify pigment cell count and body length. All points were manually added using Cell Counter in FIJI. The three images represent three of the 50 slices analyzed; points remained between each slice to prevent cells from being counted multiple times. Blue dots denoted with “1” represent pigment cells, magenta dots denoted with “4” represent preoral arms, golden dots denoted with “5” represent postoral arms, and pink dots denoted with “6” represent the apical end.

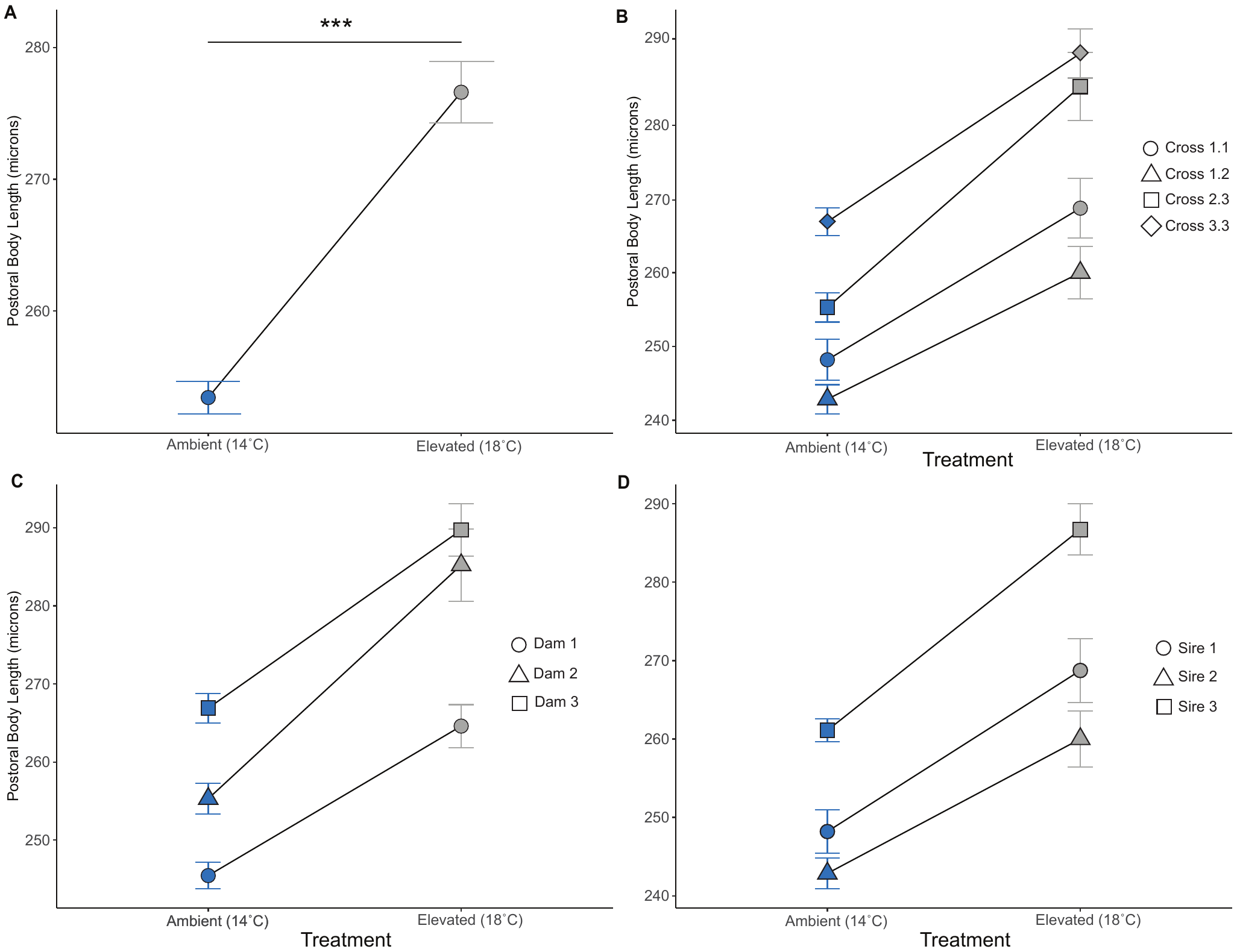

**Supplemental Figure 2: Embryos developed at elevated temperature exhibit longer pre-oral arms** A) Difference in preoral body length between larvae which developed in ambient temperatures (blue) compared to larvae which developed in elevated temperatures (grey) (*p*_lmer_=0.0211). B) Differences in preoral body length between dams. C) Preoral body length differences between genetic crosses. D) Differences in preoral body length between sires. Asterisks denote significant differences (*p*_lmer_ <0.05).
